# Pulsatile flow dynamics maintain pulmonary arterial architecture

**DOI:** 10.1101/2025.06.30.662470

**Authors:** Stephen Spurgin, Lauren Thai, Tina Wan, Christopher P. Chaney, Mitzy A. Cowdin, Suren V. Reddy, M. Tarique Hussain, Munes Fares, M. Luisa Iruela-Arispe, Thomas Carroll, Andrew D. Spearman, Ondine Cleaver

## Abstract

Single ventricle congenital heart disease (SV-CHD) is a uniformly lethal condition requiring the Glenn surgery, which as a side effect eliminates arterial pulsatility and contributes to pulmonary vascular complications. In Glenn patients, we quantified pulsatility loss in each dimension of force (flow, pressure, and stretch) using cardiac catheterization and MRI. To model and investigate the individual impact of each dimension of pulsatility loss on the pulmonary vasculature, we applied isolated pulsatile and non-pulsatile mechanical stimuli to pulmonary arterial endothelial cells (ECs) in vitro. We found that each dimension of force triggered distinct transcriptional responses, revealing force-specific regulation of structural and signaling pathways. Pulsatile stretch uniquely stimulated EC secretion of PDGFB, a key driver of vascular smooth muscle cell (vSMC) recruitment. In a rat Glenn model, loss of pulsatility led to vascular wall thinning, confirming *in vivo* relevance. Our findings uncover a mechanistic link between endothelial stretch sensing and PDGFB-mediated EC-vSMC crosstalk, essential for maintaining pulmonary artery architecture. Clinically, these insights suggest that restoring or mimicking pulsatile forces may help preserve vascular integrity and prevent remodeling in SV-CHD patients.

## Introduction

Globally, 1 in every 2000 babies is born with a heart that has a single functional ventricle (1). Surviving this congenital abnormality requires early and significant medical intervention. A single ventricle cannot perform the work of two for very long, nor can it direct an equal amount of blood flow to the lungs (low vascular resistance) and the body (high vascular resistance). For children with single ventricle congenital heart disease, survival depends on a series of remarkable surgeries: the Glenn and the Fontan. In the circulation created by these procedures, venous blood flows directly to the lungs without the support of a pumping ventricle, while the single functional ventricle supplies pulsatile flow to the rest of the body (2).

A frequent consequence of surgical replumbing of the vasculature in single ventricle patients is the development of a wide variety of vascular malformations: A) diffuse microvascular pulmonary arteriovenous malformations (PAVMs) in the Glenn stage and B) larger, tortuous collateral vessels in both Glenn and Fontan stages. It has been shown that the microvascular lesions that arise in the Glenn are reversible with restoration of the direct flow of liver blood to the lungs in the Fontan (albeit still lacking cardiac pulsatile flow). This has led to the “hepatic factor” hypothesis (3–5). However, interestingly, the proximal macrovascular collateral vessels in the Fontan do not regress (6, 7). Among the abundant potential pathogenic factors—chronic hypoxemia, chronic inflammation, and loss of direct hepatopulmonary blood flow (3, 5, 8)—a role for pulsatility has long been suggested (9–11), but never investigated at the mechanistic level.

Pulsatility, for its part, is a critical characteristic of arterial blood flow. Arteries—whether systemic, pulmonary, or umbilical—are *defined* by the presence of pulsatile blood flow, not by their color or oxygen content. The structure of the arterial wall—with thick smooth muscle layer and elastic laminae—reflects the need to withstand the high pulsations of arterial pressure and stretch. Veins, on the other hand, have less hemodynamic pulsatility and a thinner smooth muscle layer (media) (12). Critically, the histopathology of vascular malformations often reveals absent or patchy coverage of the smooth muscle layer (13, 14). This mural insulation must be broken down to form new connections between vessels. EC proliferation is a then a critical step in the pathogenesis of vascular malformations (13). Once a connection is formed, the transmission of pulsatile, arterial blood flow to venous vessels has been shown to induce “arterialization” (medial thickening, intimal hyperplasia) of the venous vessels (15, 16), further highlighting the relationship between arterial hemodynamics and the architecture of the vascular wall.

Although arteriovenous differences are most visually evident by their mural coverage, the endothelial cells of arteries and veins are themselves distinct. They have unique transcriptional and proteomic signatures (17), and unique developmental origins (18, 19). The impact of cardiac pulsatility on the arterial side of the vasculature has been appreciated in the early stages of vascularization (20)—however, the role of *pulsatility* in maintaining arterial and venous endothelial identity and function *after* birth has received very limited attention. Scientific investigation of EC mechanobiology has focused on other physical forces, such as the cellular impact of low versus high shear stress, or of laminar versus disturbed flow (21). Despite the clinical centrality of pulsatility, it remains an under-investigated aspect of vascular biology.

ECs are remarkably sensitive to the physical forces transmitted by blood flow. Endothelial mechanosensitive components include cell-cell adhesion molecules (integrins, gap-junctions, cadherins), extracellular matrix (ECM) sensing components, ion channels, and more (21, 22). The forces sensed by these molecular components are applied to Ecs in three dimensions: shear stress , pressure, and stretch (23). Each dimension of force therefore exerts distinct stresses and deformations to ECs. Laminar shear stress (LSS) is the frictional drag force that occurs along the inner (apical) surface of the vessel endothelium (24). Stretch is the effect downstream of each heartbeat, as vessels widen radially to accommodate an ejected volume of blood and their circumference expands (25). Hydrostatic pressure is generated by ventricular contraction, and the driving force behind stretch (26). Notably, ECs align parallel to the direction of LSS (blood flow) but perpendicular to the direction of stretch (27)—further highlighting the need to consider each dimension’s impact separately. All three biomechanical forces have all been shown to alter vascular ECM (28), as well as endothelial and smooth muscle transcriptional programs (29, 30). However, the unique impact of pulsatility within each dimension of force on vascular homeostasis remains a critical knowledge gap.

The endothelium and vascular smooth muscle communicate closely. Critically, cross-talk between ECs and vSMCs is mediated by EC-secretion of platelet derived growth factor B (PDGFB), which binds to PDGF receptor 0 (PDGFR 0) on vSMCs, where it drives proliferation and dictates mural cell coverage from large vessels down to the capillary level (31, 32). Loss of Notch signaling in pericytes downregulates PDGFRB in vSMCs in mouse models, leading to AVMs (33). Conversely, deleting PDGFB specifically from ECs has shown remarkable dilation and arteriovenous shunting in the lungs of mice (34), and loss of PDGFB has been observed in human vascular malformations (35). The question arises as to whether different hemodynamic forces coordinate the maintenance of vascular structure via EC responses to these forces.

Here, we investigate the biomechanical forces experienced by pulmonary ECs in Glenn patients and identify pulsatility of blood flow as critical to maintenance of the lung vasculature. We first use combined cardiac catheterization (cath) and cardiac magnetic resonance imaging (MRI) to quantify and compare the pulsatility of force that is applied to pulmonary arterial ECs in children with normal cardiopulmonary anatomy and in those with Glenn circulation. After noting a clear loss of pulsatility in the Glenn pulmonary arteries, we model those changes using cultured human pulmonary arterial endothelial cells (HPAECs) to define the specific transcriptional programs driven by each individual dimension of force. We demonstrate that pulsatile stretch in vitro, which mimics expansion of the arterial vessel intima during cardiac ventricular systole, significantly impacts the transcriptional landscape of ECs. Specifically, we show that pulsatile stretch stimulates ECs to secrete PDGFB. In addition, using our previously published in vivo rat model of the Glenn surgery, where pulmonary arteries (PAs) experience loss of pulsatile stretch, we demonstrate significant reduction of PDGFB in PA ECs. Finally, we build on our recent work in a rat model of the Glenn surgery(36), which showed early and progressive arteriovenous shunting, to demonstrate significant thinning of pulmonary arterial vascular walls.

Together, our study identifies pulsatile stretch as a critical and underappreciated dimension of arterial force that drives EC-secretion of PDGFB, promoting vascular smooth muscle endowment and providing needed structural support for arterial vessels. Gaining insight into the impact of *pulsatility* of hemodynamic force is a critical and novel approach to understanding clinically relevant endothelial cell biology.

## Results

### The Glenn shunt directs venous blood flow to pulmonary arteries

Arterial vessels normally experience blood flow directly from a pumping cardiac ventricle, adding pulsatility to the magnitude of hemodynamic forces (**Figure 1A**). By reviewing the known characteristics of three distinct pairings of arteries and veins (systemic, pulmonary, and umbilical), we note that pulsatile flow is a shared, definitional quality of arteries (**Supplemental Figure 1**). Pulsatility is thus an inherent quality of blood flow in arterial vessels. After a Glenn surgery, however, the pulmonary arteries no longer receive blood flow directly from a pumping heart ventricle. Instead, pulmonary blood flow is provided by the Glenn shunt: direct venous flow through the superior vena cava (SVC), which is anastomosed to the pulmonary arteries (PA) (**Figure 1A’**). ECs that make up the inner lining of pulmonary arteries downstream of normal cardiac vessel anatomy are subjected to force in three dimensions: flow/sheer stress, pressure, and circumferential stretch (**Figure 1B**) (23). Here, we assess these forces in the Glenn lung arteries.

**Figure 1.**
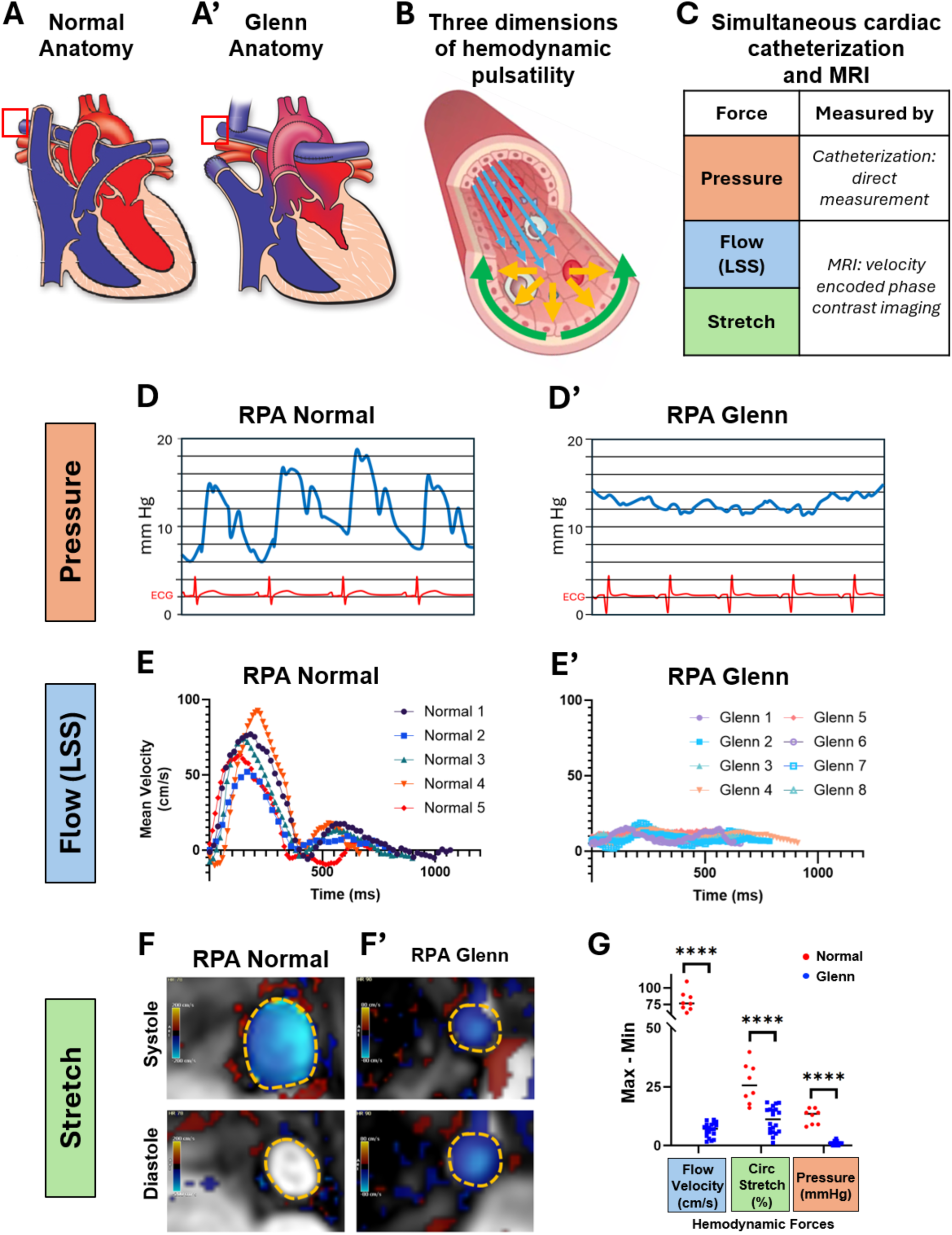
Loss of pulsatility in three dimensions within Glenn pulmonary arteries. (A, A’) Diagrams of representative normal and Glenn cardiopulmonary anatomy. Red box highlights location of force pulsatility measurements. **(B)** Hemodynamic forces act on ECs in three dimensions: shear/flow (blue), pressure (orange), and circumferential stretch (green). (Schematic adapted from graphic on Ibidi.com). **(C)** Outline of data collection method for each dimension of hemodynamic force. **(D, D’)** Example pressure waveforms from the right pulmonary artery of normal and Glenn patients. Electrocardiogram shown in red for correlation to cardiac cycle. RPA, right pulmonary artery. **(E, E’)** Variation in flow velocity of the proximal right pulmonary artery measured by cardiac MRI throughout a cardiac cycle from systole (start) to end-diastole (end). Variability in length of time for each patient reflects the variable heart rate within each subject. **(F, F’)** Cardiac MRI velocity encoded phase contrast images with color showing velocity of blood flow out of the plane of the image slice. Dotted orange line shows RPA border. RPA diameter increases during systole in anormal patients but does not increase during systole in Glenn patient. **(G)** Quantification of pulse difference of each dimension of force, obtained during combined cardiac catheterization and cardiac MRI of normal (n=20) and Glenn (n=20) patients, demonstrating loss of pulsatility in the Glenn. Circumference % change is calculated from baseline in diastole.

### Pulsatility loss in three dimensions in Glenn patients

To quantitatively determine the loss of pulmonary arterial pulsatility in patients with Glenn anatomy, we used combined cardiac catheterization and cardiac MRI (cath/MRI) in 20 Glenn patients to obtain simultaneous data in all three dimensions of force: pressure, flow, and circumferential stretch (**Figure 1B**). These data were compared to data obtained in 20 age- and gender-matched patients with normal cardiopulmonary connections (see patient demographics in **Supplemental Table 1**). First, as has long been observed clinically, we observed pulsatility loss in the pulmonary arterial *pressure* of Glenn patients (**Figure 1C, C’**). Similarly, we noted that the Glenn shunt leads to non-pulsatile shear stress in the pulmonary arteries, as seen by a comparison of the pulmonary artery velocity profiles in normal (**Figure 1D**) and Glenn (**Figure 1D’**) patients throughout the cardiac cycle. Finally, we found that the circumference of proximal right pulmonary artery does not expand repeatedly with each heartbeat in Glenn patients, demonstrating loss of pulsatile *circumferential stretch* (**Figure 1E,E’**).

### Unique transcriptome for each dimension of pulsatile force

ECs are highly responsive to mechanical force, and can react to differences in both magnitude and direction of force.(37) Given the observed loss of pulsatility in Glenn patients, we next investigated the relative impact of pulsatility within each dimension of force on ECs.

Using our patient cath/MRI pulsatility data, we designed in vitro experiments that model the pulsatility loss defined in Glenn patients. Each isolated, single dimension of force was applied to cultured primary human pulmonary artery endothelial cells (HPAECs) in a pulsatile and non-pulsatile manner. For pulsatile shear stress, we applied 0-15 dyn/cm^2^ shear stress at 1 Hz, which mimics the frequency of a resting heart rate. For pulsatile pressure, we modified the Ibidi equipment to expose HPAECs to columns of media positioned to alternately provide 5 or 25 mmHg pressure, at 1 Hz, in the absence of flow or stretch. For stretch, we used a Flexcell system to stretch HPAEC by 10% of their baseline in a single axis (again at a frequency of 1 Hz). To assess stable transcriptional changes between non-pulsatile and pulsatile conditions and thus improve in vivo relevance of our findings, HPAECs were exposed to 48 hours of force or pulsatile force (**Figure 2A**). To best mimic in vivo conditions in the years of the Glenn stage, pulsatile stretch was compared to no stretch rather than continuous stretch, which does not happen in vivo. We observed similar physical changes as have been reported previously: ECs align parallel to the axis of flow and perpendicular to the axis of stretch and exhibit no alignment preference under continuous or pulsatile pressure (**Figure 2B**).

**Figure 2.**
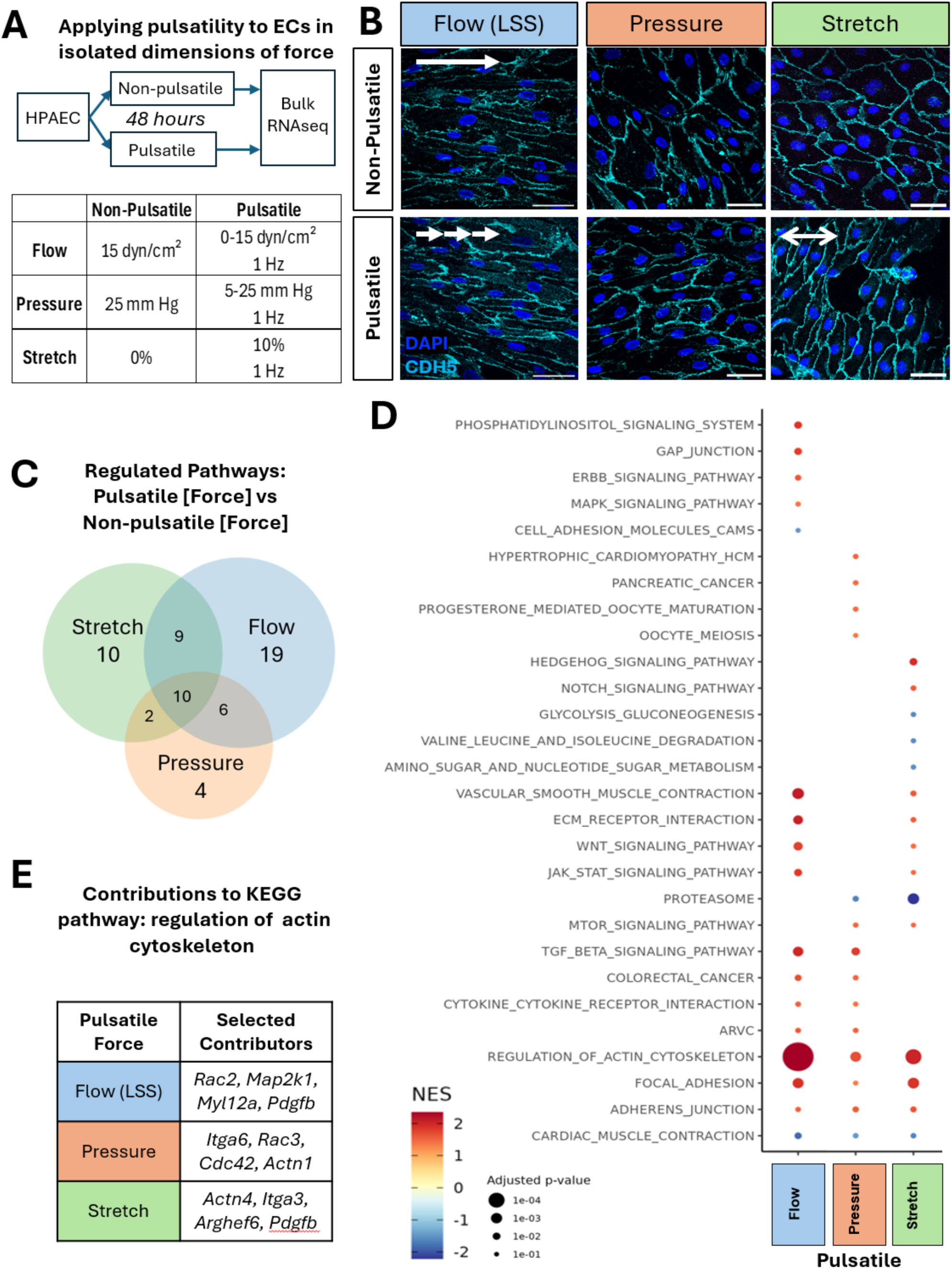
Pulsatility drives both similar and distinct signaling pathways within each dimension of force. **(A)** Experimental conditions: HPAECs were exposed to 48 hours of a single dimension of either non-pulsatile or pulsatile force, and bulk RNA sequencing was performed. **(B)** GSEA of bulk RNAseq data shows overlapping and unique pathways activated by each dimension of force. **(C)** Representative images of the cell-shape adjustments that occur in response to the different forces are shown. **(D)** Normalized enrichment score (NES) and adjusted p-value (adj p-val) shown for select signaling pathways shows greater impact of pulsatility within flow and stretch relative to pulsatility of pressure. **(E)** Key genes with the actin cytoskeleton pathway that are upregulated by each dimension of force. **(F)** *Pdgfb* was downregulated in 11 of dysregulated pathways identified in E.

The unique changes in cell shape in response to each axis of force highlight the need to examine them individually. We first investigated the overall pathways that were regulated by pulsatility. By Gene Set Enrichment Analysis (GSEA) of bulk RNAseq data, we noted that each dimension of force drove changes in unique signaling pathways, and that the impact of shear stress was the most distinct. Overall, in HPAECs, laminar flow affected 19 unique pathways, stretch 10 unique pathways, and pressure 4 unique pathways (**Figure 2C**). A wide range of pathway types were regulated, many related to changes in structural cellular components, growth, or interaction of the cell with ECM (**Figure 2D**). We note that mitogen-activated protein kinase (*Mapk*) and epidermal growth factor receptor (*ErbB*, also known as *HER*) signaling were only induced by pulsatility of flow, while Hedgehog and Notch signaling were only induced by pulsatility of stretch. Pulsatile pressure did not uniquely regulate any obviously EC-related pathways, but did upregulate the hypertrophic cardiomyopathy pathway.

In addition to these dimension-specific pathways, we find other pathways that are influenced across multiple dimensions of force. We found 10 pathways upregulated by all 3 dimensions of force, and 17 pathways induced by two of the three dimensions. We note that shear stress, stretch, and pressure all drove changes in the pathway “Regulation of Actin Cytoskeleton.” However, an analysis of the leading-edge subset (the genes within the pathway that are most significantly changed and thus responsible for the overall significance of the pathway) revealed upregulation of different genes by each dimension of force. For example, within the “Regulation of Actin Cytoskeleton” pathway, pulsatile pressure shows *Actn1* and pulsatile flow yields *Map2k1*, while pulsatile stretch promotes *Pdgfb*, *Arghef6,* and others (**Figure 2E**). Different dimensions of force therefore drive different transcriptional changes that converge on similar pathways.

### Pulsatility does not alter arteriovenous genes in vitro

To test the impact of each dimension of force on ECs, we next analyzed gene transcription. Using the culture platforms shown in **Figure 3A**, we assessed bulk RNAseq after application of single dimensions of pulsatile and non-pulsatile force. Strikingly, we found very little overlap between genes enriched in ECs under different dimensions of force, suggesting distinct responses to distinct forces. Importantly, we did not find significant changes in baseline expression of common EC genes between the three experimental set-ups (**Supplemental Figure 2**), indicating no basic alteration of endothelial fate. Of the genes significantly regulated by pulsatility, 93.4% (1205/1290) were driven by a single, unique dimension of force. Pulsatility of stretch accounted for over two-thirds (888 genes, 68.9%) of these genes (**Figure 3B**). Only 59 genes were uniquely induced by pulsatility of laminar flow, while 343 were uniquely induced by pulsatility of pressure. We found less overlap in individual genes regulated by pulsatility than there was in overall pathways. Twenty-two genes were regulated by pulsatility of both LSS and stretch, 56 genes by stretch and pressure, and 5 by pressure and LSS.

**Figure 3.**
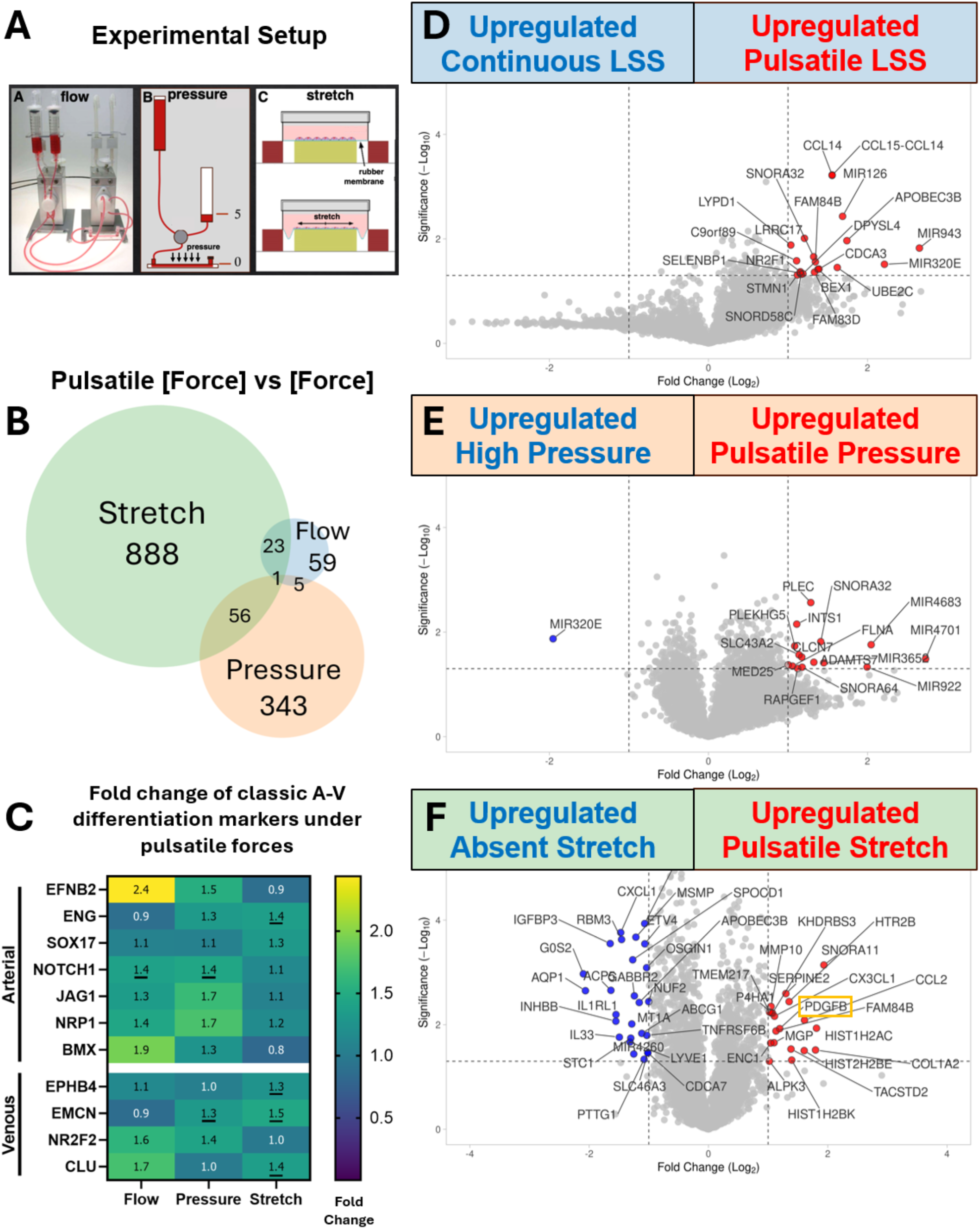
Impact of pulsatile stretch is greater than pulsatile pressure or flow but does not change classical arteriovenous differentiation. **(A)** Diagram of cell culture setup to apply isolated dimensions of force to ECs. **(B)** Venn diagram of the number of genes significantly up- or down-regulated by pulsatility of flow, pressure, or stretch. **(C)** Fold change (pulsatile/non-pulsatile) of any classic EC arteriovenous differentiation markers by bulk RNAseq. Color scale denotes fold change, exact number in each box. Significant changes (p-values < 0.05) are underlined. **(D-F)** Volcano plots of significantly regulated genes within each dimension of pulsatile force: **(D)** LSS; **(E)** pressure; and **(F)** stretch.

Finally, we hypothesized that application of pulsatile force on cultured ECs would induce greater expression of genes previously shown to be specific to arteries (17). We asked whether the classical markers of arteriovenous (AV) EC differentiation were found in these groups of genes. In cultured HPAECs, we found no dimension of pulsatility that significantly affected the expression of a majority of developmental arterial (*Efnb2*, *Eng*, *Sox17*, *Notch1, Jag1, Nrp1, Bmx)* or venous (*Ephb4*, *Emcn*, *Nr2f1*, *Nr2f2*/*Coup-TFII, Clu*) driver genes (**Figure 3C**). Similarly, we found no significant changes arterial or venous standard marker genes in rat Glenn arterial ECs compared to sham operated rats (data not shown).

We next asked if there were genes specifically induced by pulsatile LSS, pressure or stretch, when compared to the non-pulsative forces. After applying a p-value cut-off of 0.05 and fold change cut-off of 2, we found that pulsatile stretch drove a change in 44 genes, compared to 20 genes by pulsatility of flow, and 16 genes by pulsatility of pressure (**Figure 3D-F**). Expression of these hits in data from other dimensions of force is shown in **Supplemental Figure 3**. Genes upregulated under pulsatile LSS included many factors involved in cell proliferation and immunity (red highlighted genes in **Figure 3D**), and genes upregulated under pulsatile pressure included factors associated with the cytoskeleton and RNA processing (red highlighted genes in **Figure 3E**). Gene expression was particularly impacted by application of pulsatile stretch, with many factors involved in tissue remodeling and inflammation being upregulated (red highlighted genes in **Figure 3F**) and factors driving cell division and cell survival pathways (blue highlighted genes in **Figure 3F**).

### Identification of *Pdgfb* as a stretch-induced gene in pulmonary arterial endothelium

As stretch accounted for the majority of pulsatility-regulated genes, we utilized published single cell RNA sequencing of human lung tissue (LungMAP(38, 39)) to assess the cell type-specific expression pattern of candidate genes. We reasoned that narrowing the investigation to EC-specific genes might identify endothelial factors that sense flow and normally respond by bolstering matrix or cellular components of the arterial media that would increase resistance in proportion to hemodynamic load. *Mgp*, *Cx3cl1*, and *Pdgfb* were highly expressed in a good proportion of ECs under stretch, but we noted high EC specificity of the latter two genes (**Figure 4A**). Analysis of the expression of these ligands and their receptors (*Cx3cl1R* and *Pdgfrb*, respectively) underscored the pulsatility-induced crosstalk between ECs and immune cells (for *Cx3cl1*) and between ECs and perivascular cells (for *Pdgfb*) (**Figure 4B-D**). As PDGFB is i) well known to promote smooth muscle cell proliferation and recruitment (31), ii) a key component of the GSEA for ECs under stretch (**Figure 2**), and iii) a significant hit in their differential gene expression (**Figure 3**), we investigated further its role in pulmonary vessels.

**Figure 4.**
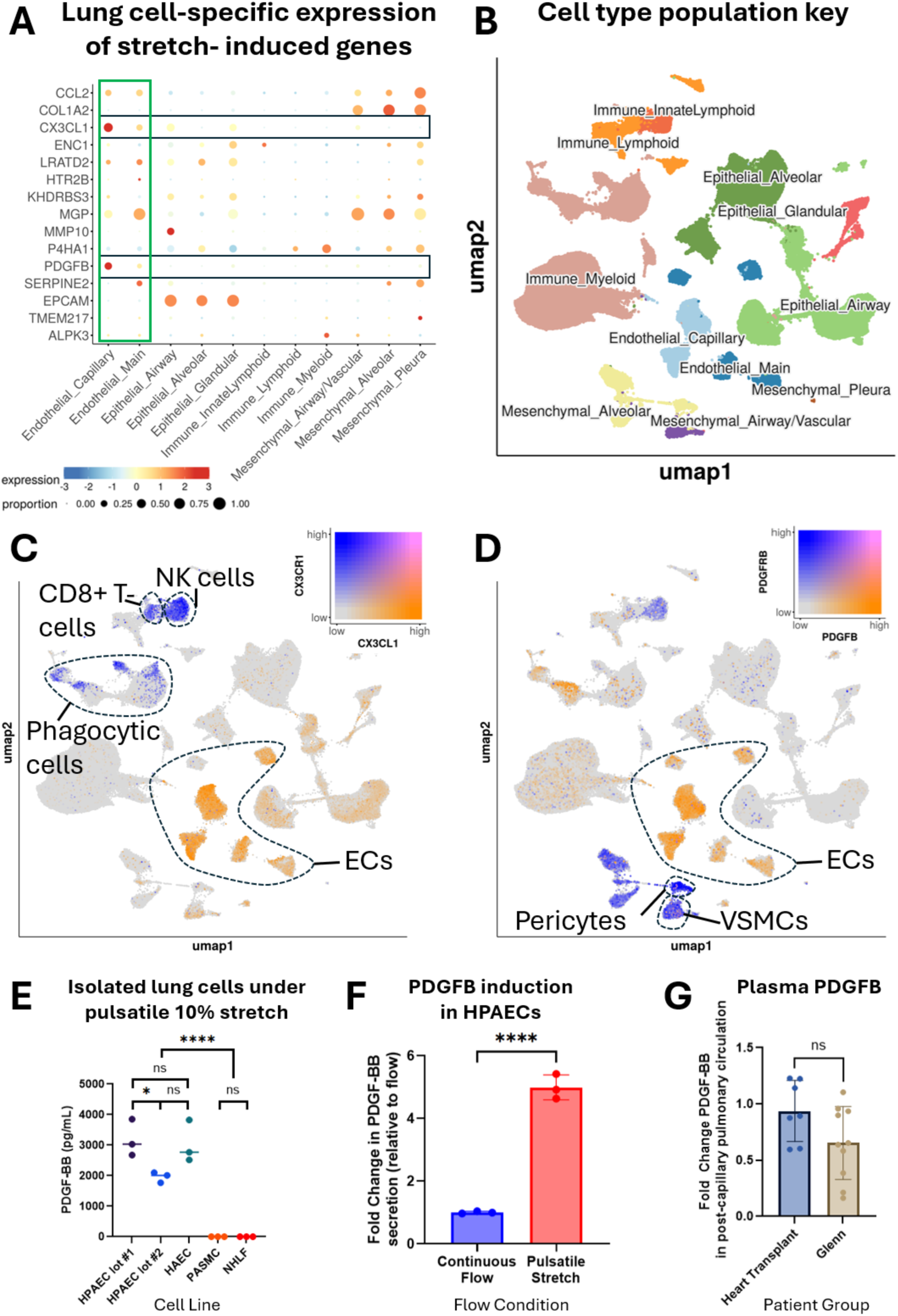
PDGFB expression in human lungs is endothelial-specific and stretch-induced. **(A)** Genes upregulated by pulsatile stretch in ECs were analyzed within the LungMAP single cell RNA sequencing data of human lung tissue (347,970 cells). Expression within ECs (green box) was compared to other cell types. CX3CL1 and PDGFB (black boxes) were uniquely expressed by ECs. **(B)** Cell population groupings as annotated by LungMAP cell x gene portal. **(C)** Gene co-expression analysis shows expression of *Cx3cl1* in ECs with its receptor *Cx3cr1* in immune cells. **(D)** Expression of *Pdgfb* is high in ECs, with its receptor *Pdgfrb* found in pericytes and vascular smooth muscle. **(E)** Different cell lines comprising the primary components of a blood vessel (endothelium, smooth muscle, fibroblasts) were exposed to pulsatile uniaxial 10% stretch at 1 Hz and only ECs secreted PDGFB into the media (as measured by ELISA). **(F)** Pulsatile stretch of cultured HPAECs induces 5x more PDGFB secretion than continuous flow. Error bar shows 1 s.d. **(G)** PDGFB measured by ELISA from patient plasma does not show a significant arteriovenous concentration gradient or difference between normal and Glenn patients. Error bar shows 1 s.d.

### Pulsatile stretch uniquely drives EC PDGFB secretion

We next asked if stretch-induction of PDGFB expression was specific to ECs or also present in other cellular components of the vessel wall (vascular smooth muscle or fibroblasts). We applied cyclic 10% stretch to HPAECs from two different patients, human aortic endothelial cells (HAECs), human pulmonary artery smooth muscle cells (PASMCs), and normal human lung fibroblasts (NHLFs). We found that pulsatile stretch robustly induced secretion of PDGFB exclusively from endothelial cell lines (**Figure 4E**). As PDGFB is known to be induced by flow (40), we compared PDGFB secretion from ECs under continuous flow (mimicking venous condition) to pulsatile stretch (mimicking arterial condition). Induction of PDGFB by pulsatile stretch was 5x higher than the induction by laminar, continuous flow (**Figure 4F**).

We next assessed whether we could detect EC secretion of PDGFB in human patients following similar pulsatile stretch-induction. We anticipated that we would not be able to detect a change in the circulating blood, as PDGFB is secreted basally to the perivascular space, not apically into the blood stream, as shown previously (41). To do this, we tested the blood of patients with either normal or Glenn anatomy. As expected, analysis of PDGFB expression in plasma obtained from the pulmonary artery and pulmonary vein revealed significant variation in local PDGFB expression in both normal and Glenn patients that could not be correlated with pulmonary arterial pulsatility (**Figure 4G**).

### Decreased PDGFB expression in pulmonary arteriole ECs of the Glenn rat model

Given the finding that stretch induces PDGFB secretion from pulmonary ECs, we utilized our animal model of the Glenn surgery to test whether arteries receiving non-pulsatile flow exhibited decreased endothelial PDGFB synthesis in vivo. We performed an end-to-end anastomosis of the left SVC (L-SVC) and the left pulmonary artery (LPA) (**Figure 5A, A’**). This mimics the Glenn surgery, directing only non-pulsatile venous blood flow from the head and upper extremity into the pulmonary artery. As previously described, these rats develop hypoxemia secondary to intrapulmonary arteriovenous shunting.(36) The hypoxemia develops rapidly (after two weeks) and is progressive (to six months (36)).

**Figure 5.**
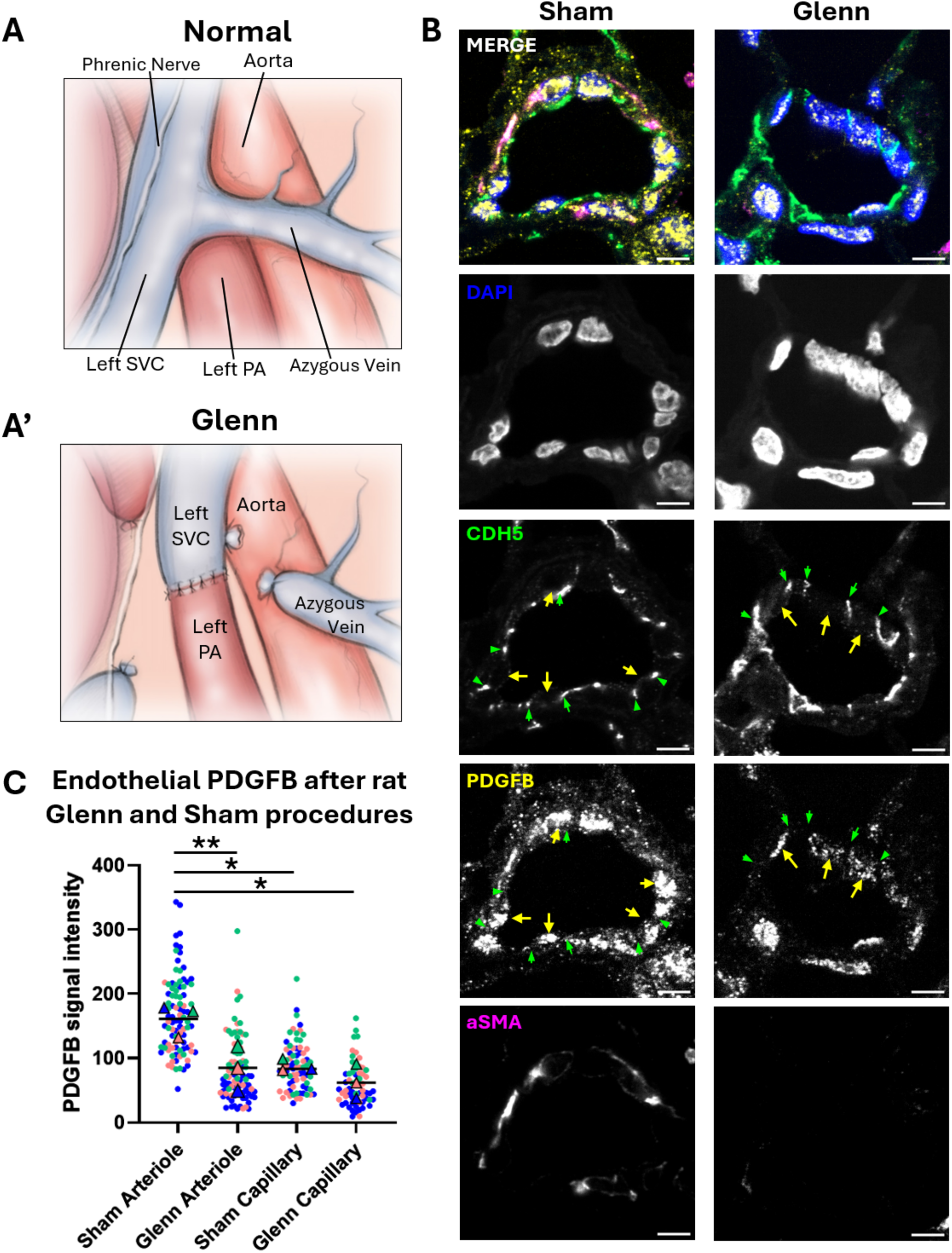
Loss of PDGFB in pulmonary endothelium of rats after Glenn surgery. **(A)** Schematic of normal rat vascular anatomy and (**A’**) rat Glenn surgery (adapted from Wan 2024 with permission). **(B)** Representative immunofluorescent (IF) staining of pulmonary arterioles from left lung of sham and Glenn-operated rats. Pulmonary arterioles were identified by presence of VECAD (cyan) and SMA (pink), near lung periphery, and less than 50 µm in diameter. Scale bar is 5 µm. **(C)** Quantification of PDGFB staining in pulmonary arteriolar and capillary ECs in sham (N=91 arteriolar ECs, N=74 capillary ECs) and Glenn-operated (N=89 arteriolar ECs, N=62 capillary ECs) rats demonstrates significant loss of EC-derived PDGFB in the Glenn lung. 5 vessels each from the left lung of 3 sham and 3 Glenn rats were analyzed.

To test whether arteries receiving non-pulsatile flow exhibited decreased PDGFB synthesis in their ECs, we focused on the distal lung arterioles—in the vicinity of where AVMs form in our patients, and where EC-specific expression of PDGFB is highest by single cell RNAseq (**Figure 4A**). To localize these arterioles and the PDGFB expression within the ECs, we used immunofluorescent staining with antibodies recognizing the endothelial junctional marker VE-cadherin (CDH5), the vascular smooth muscle marker alpha smooth muscle actin (aSMA) and the PDGFB protein (42). We found significant reduction of endothelial PDGFB in arterioles of the Glenn rat (**Figure 5B**). Critically, we also noted loss of smooth muscle surrounding these distal arterial vessels in the Glenn. Quantification of PDGFB IF signal from 5 arterioles each from 3 sham and 3 Glenn-operated left lungs revealed a significant decrease in endothelial PDGFB expression in the Glenn arterioles, but not in alveolar capillary ECs (**Figure 5C**).

### Rat Glenn displays reduced pulmonary arterial smooth muscle

We hypothesized that a further consequence of reduced PDGFB in pulmonary ECs under the non-pulsatile Glenn hemodynamics would be loss of the smooth muscle layer. We analyzed sagittal sections of the left lung of 9 Glenn rats and 5 sham-operated rats and performed immunohistochemistry (IHC) for smooth muscle actin (SMA). Strikingly, we found a significant decrease in arterial smooth muscle thickness (SMA+ cells) (**Figure 6A, B**), while venous smooth muscle thickness remained unchanged (**Figure 6C**). We observed this reduction in vascular wall thickness in both the largest proximal arteries (**Figure 6A-C**), as well as the distal arterioles, less than 0.1 mm in diameter (**Figure 6D-G**).

**Figure 6.**
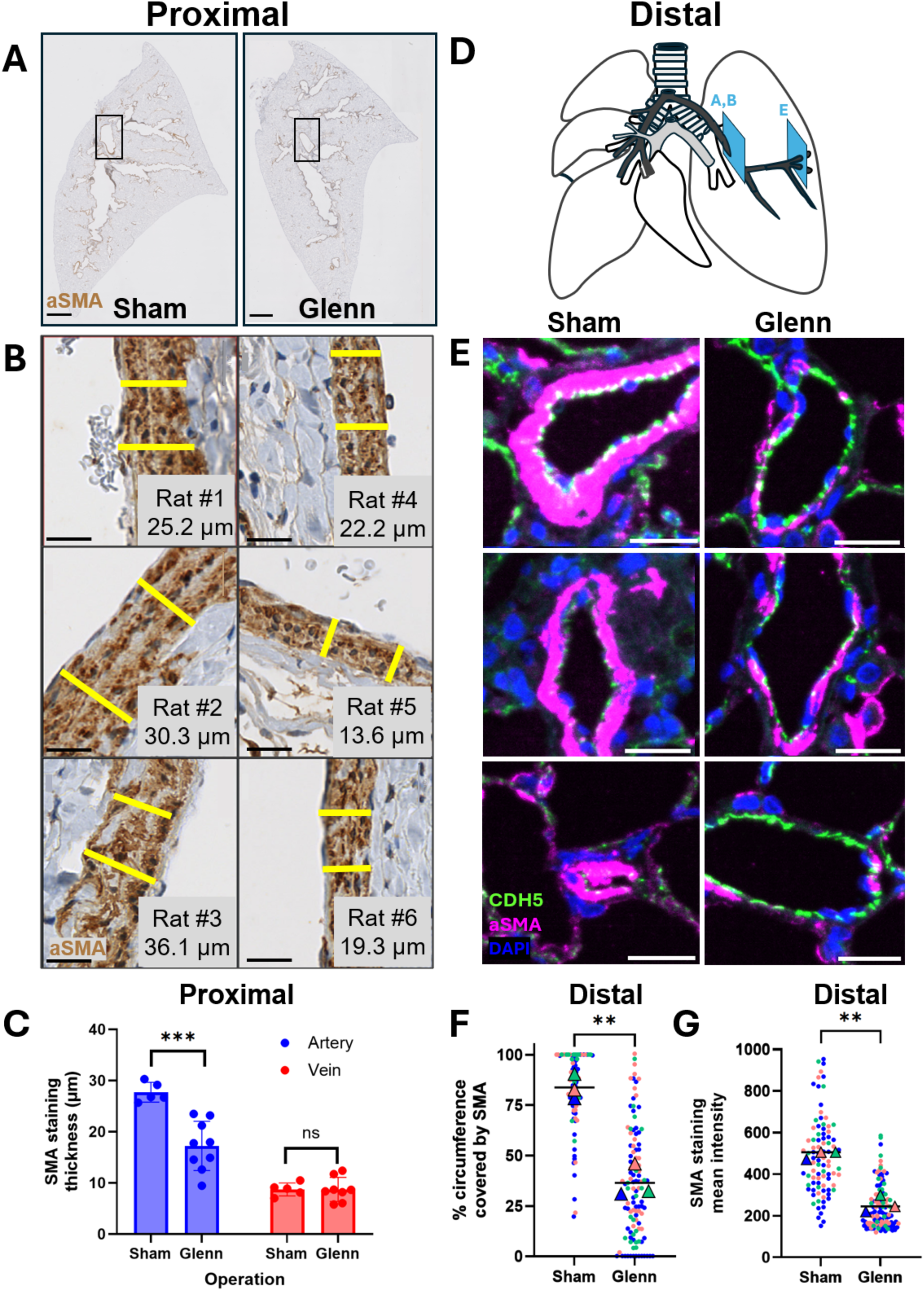
Loss of smooth muscle layer in pulmonary arteries of rats after Glenn surgery. **(A)** Example sagittal sections of lung tissue from sham and Glenn operated rats, stained by IHC for smooth muscle actin (SMA). Black boxes show typical perihilar location of analysis of the most proximal portions of pulmonary arteries examined. Scale bar = 2 mm. **(B)** Left panels - Example images of SMA+ media thickness from 3 different sham operated rats. Scale bar = 20 µm. Right panels - Example images of SMA+ media thickness from 3 different Glenn rats. Scale bar = 20 µm. **(C)** Quantification of SMA+ tunica media from proximal arteries versus veins. **(D)** Schematic showing approximate location of pulmonary vessels assayed in A, B and E. **(E)** Examples of images of SMA+ media thickness in distal arterioles from 3 different sham (left panels) and Glenn (right panels). Distal lung vessels < 0.1 mm were identified using IF staining for CDH5 (green) in Sham (n=3 rats, n=42 vessels) and Glenn (n=3 rats, n=56 vessels) rat lungs, and staining of SMA (pink) was used to quantify the circumferential coverage **(F)** and intensity **(G) of** SMA surrounding these arterioles.

## Discussion

In this study, we identify hemodynamic pulsatility as critical to maintenance of the pulmonary arterial architecture (**Figure 7**). We show that in patients with Glenn circulation the shunted pulmonary blood flow—from the SVC directly to the PA—exerts no pulsatile stretch on the pulmonary artery. Using transcriptomic analysis, we show that cultured pulmonary arterial ECs exposed to each dimension of force elicit distinct molecular signatures. We find that pulsatile stretch induces significant endothelial secretion of PDGFB, known to mediate recruitment of vSMCs to blood vessels (31, 32). As lung biopsies from living patients were not obtainable, our rat model of the Glenn circulation allowed us to investigate the effect of pulsatility loss in vivo. Using this animal model, we show that non-pulsatile, venous flow within an arterial vessel, results in loss of PDGFB protein from ECs and concomitant thinning of the vascular mural wall. Our complementary in vitro and in vivo findings identify a new target signaling pathway for novel therapeutic approaches to treat the diverse vascular malformations that occur in Glenn patients.

**Figure 7.**
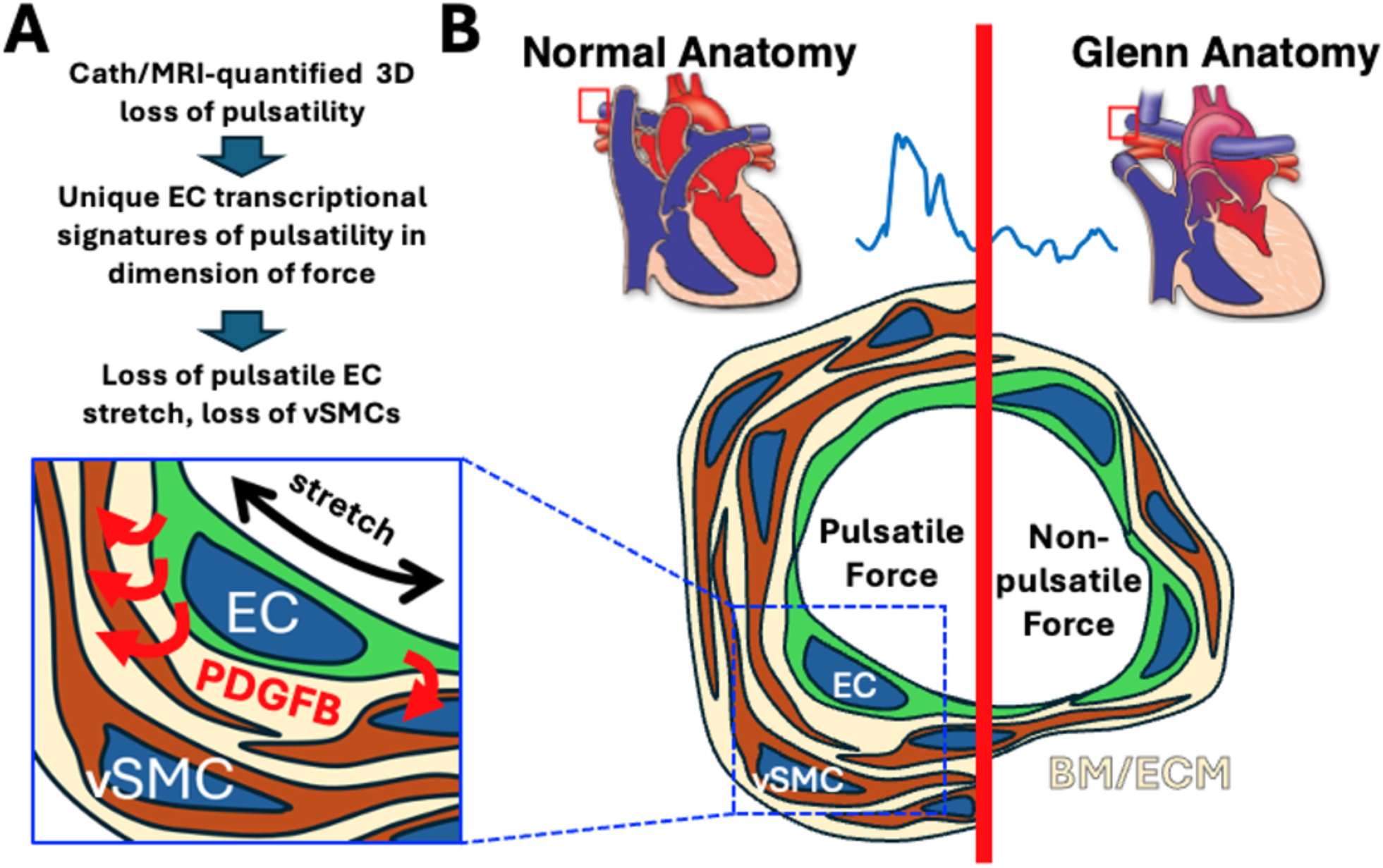
Outline of primary findings. **(A)** Overview of experimental results. **(B)** Comparison of findings between pulsatile and non-pulsatile conditions. Red boxes indicate region of pulmonary arteries analyzed in patients with either normal lung vascular architecture or Glenn anatomy. Blue box area shows increased PDGFB secretion from ECs following blood circulation-induced stretch. EC, endothelial cell (green); vSMC, vascular smooth muscle cell (brown); BM, basement membrane, and ECM; extracellular matrix (beige).

Despite clinical appreciation of the fact that Glenn patients experience venous-type, non-pulsatile blood flow in pulmonary arteries, and even though pulsatility loss was the first hypothesized mechanism for Glenn PAVMs (43, 44), pulsatility loss has never been formally documented in three dimensions. Until now, studies comparing surgical outcomes of Glenn patients have only occasionally been able to report some dimensions of pulsatility loss (9, 45), although most studies report none (46–52). This clinical knowledge gap has limited our understanding of the impact of pulsatility within these patients. Our use of combined cardiac catheterization and cardiac MRI (cath/MRI) ensures that these interdependent forces are measured simultaneously, providing with high temporal accuracy the first complete 3D view of pulsatility loss in the Glenn pulmonary arteries.

Our pulsatility metrics merit two points of discussion. First, we utilize flow velocities as a surrogate for wall shear stress (WSS)—the direct hemodynamic force applied to the endothelium. As WSS depends directly on the velocity gradient, velocity is a reasonable approximation for this investigation. Given the limited availability and accuracy of WSS measurements with cardiac MRI, use of velocity instead of the highly regionally variable WSS supports an easier and more rapid clinical application. Second, the relationship between pressure and stretch (vascular compliance) is not directly tested in our study. This will be a critical area of investigation in the future, to answer the question of how much pulsatility (of flow, pressure, or stretch) is present in the distal lung.

Indeed, estimating the magnitude of pulsatile forces across the pulmonary vascular tree is difficult. Arterial pulsatility in normal patients is likely significantly diminished as it reaches distal capillaries. Dynamic measurement of distal pulsatility of stretch in sub-millimeter vessels in vivo is technically challenging and could not be directly assessed by our methods. In a normal compliant vessel, proximal stretch acts as a capacitor, absorbing and lessening distal pulsatility—a phenomenon termed the ‘windkessel effect’ (53). This model suggests a reduction of pulsatility in the normal distal microvasculature. Our proposed mechanism—that pulsatile stretch of ECs stimulates PDGFB followed by smooth muscle coverage—is supported by the observation that normal arterial smooth muscle thins as vessels branch further into the lung (12). PDGFB was originally described as flow responsive (40), and later as stretch responsive (54), although in our in vitro work stretch elicited 5-fold more PDGFB secretion than flow. The opportunity now arises to characterize the relative contribution of shear and stretch to PDGFB secretion in these distal vessels.

We also recognize that the presence of respiratory-driven alveolar stretch (at a much lower frequency) may also contribute to distal pulsatility (55, 56). In all, these studies underscore the importance of pulsatility throughout the entire arterial network and bring up new questions about the impact of biomechanical cues on the lung vasculature.

How do ECs sense hemodynamic forces in lung vessels? We previously hypothesized that endothelial cilia could be involved in sensing pulsatility of flow (57) but disproved this possibility in the mammalian lung as pulmonary ECs are devoid of primary cilia (58). Endothelial mechanosensitive components have been well described and are present on lung ECs. Notch, Cdh5, and many other cell-surface proteins, ECM, and cytoskeletal elements are all required for proper sensing of flow or stretch (23). In particular, the protein complex of PECAM1, CDH5, VEGFR2/3, and PLXND1 plays a critical role in force-sensing, while proteins like LPHN2 and others coordinate downstream responses (59). In our data, we see significant upregulation of *Plxnd1*, as well as near two-fold increases in *Cdh5, Kdr,* and *Lphn2* (data not shown) under pulsatile pressure conditions, but not pulsatile stretch or LSS —suggesting a possible role for pulsatile pressure in priming the endothelium to respond to force. Our study thus highlights the need to consider each dimension of force separately, as we continue to evaluate the role that pulsatility plays in the maintenance of the normal vascular architecture.

One key question is how loss of pulsatility impacts arteriovenous fate in Glenn vessels. Our studies show that the Glenn shunt supplies venous-type hemodynamic forces to an artery, as blood flow pulsatility is lost following surgical intervention. The observed outcome of this change in biomechanical forces is that the artery becomes structurally more like a vein, in that its vascular wall is significantly diminished in thickness. While enhancement of arterial EC markers by pulsatility of LSS has been observed previously in vitro (20), in our studies, we did not find enhancement of classic markers of arterial EC identity under pulsatile conditions (in any dimension of force), nor did we find shifts in arteriovenous identity in our rat Glenn model (data not shown). It is possible—if unlikely—that this difference is due to the highly laminar flow provided by the Ibidi system in our work, compared to the more turbulent flow in the cone-disc viscometer used in the prior study. Separately, microvessels exposed to arterial anastomosis have been observed to increase vSMC coverage (60), however changes in EC arteriovenous identity were not noted. These observations support our conclusion that pulsatility drives the arterial vascular structure as a whole (EC + surrounding mural cells), rather than the shifting the expression of developmentally-regulated markers of arteriovenous EC identity.

Growing evidence has pointed to a potential venous origin of AVMs (61, 62), while our study focused on upstream impacts within lung arteries and upregulation of endothelial PDGFB. Endothelial deletion of *Pdgfb* (using Cdh5-CreERT2) in mouse leads to vascular malformations in mouse lungs, while arterial-specific ablation (using Bmx-Cre) does not (34). However, we note that in these studies, ablation was induced over a short timeframe (1 week) and only retinal arteries were assessed. It will be important to examine how arterial loss of *Pdgfb* over a longer timeframe, like that experienced by patients and Glenn rats, impacts downstream PAVMs. Hence, the mechanisms and time course of how upstream biomechanical cues impact arterial ECs and the downstream endothelium of capillaries and veins, and whether there are organotypic effects, remains to be seen.

Other cells of the vascular wall, such as vSMCs, also certainly sense biomechanical forces. VSMCs experience stretch similar to ECs, but not LSS as they are not in direct contact with blood flow. Interestingly, vSMCs respond differently to stretch than ECs, aligning perpendicular to stretch in vivo, rather than parallel like ECs (63). In our study and in publicly available single cell RNA sequencing from human lung, we also find that smooth muscle cells do not secrete PDGFB. We note that vSMCs from rat Glenn do not show changes in standard vSMC genes such as *myh11, tagln, myocd, cnn1*, and *pdgfrb* (data not shown). This does not rule out, however, the possible contribution of vSMCs to the ECM, especially regarding the alterations in pulmonary vascular resistance and vessel stiffening that so critically drive patient outcomes. Future studies will be necessary to investigate the responses of vascular smooth muscle to pulsatile stretch.

While we focused here on *Pdgfb*, we also note that under pulsatile flow there is a significant upregulation of *Col1a2,* a major component of vascular wall. Loss of *Col1a2* is associated with vasculopathy and aortic dilation (64), suggesting that pulsatility of stretch likely drives other EC contributions to the structure of the arterial wall—and arterial vascular identity—beyond the recruitment of vSMCs. Regulation of the ECM is a logical complimentary function of pulsatility, as the scaffold for the vSMCs and the internal elastic lamina of the largest arteries are unique features of overall arterial structure. Our work thus highlights a distinction between the traditional markers of arteriovenous endothelial identity (EC-centric) and arteriovenous vascular identity (the overall structure of the vessels cellular layers and ECM).

In addition, we also noted the upregulation of CX3CL1 in pulsatile-stretched HPAECs. CX3CL1 is a well-known chemokine that increases leukocyte recruitment and adhesion to ECs (65, 66). Multiple other chemokines (CCL2, CCL14) are regulated by pulsatility in our in vitro system. The role of the immune system in the vascular complications of Glenn circulation is a wholly unappreciated factor; however, previous work in hereditary AVMs identified accumulation of leukocytes in brain AVMs and skin telangiectasias (67–69). Collectively, these findings suggest that pulsatile stretch may impact EC-leukocyte crosstalk in pathogenesis of vascular malformations.

Translation of our findings will depend on the relative role of pulsatility in the pathogenesis of the different vascular malformations that arise in the Glenn lung. They are highly diverse, and include both large “collateral” vessel formation proximally, as well as microvascular arteriovenous malformations distally (7). As patients can present with primarily proximal, primarily distal, or both types of malformations, it is highly likely that the pathogenesis of these lesions is multifactorial, with both common and distinct elements. As previously noted, there is 50 years of evidence that a “hepatic factor” is the primary pathogenic factor for distal microvascular PAVMs. However, we suggest it is not likely the only factor. An argument for the hepatic factor as the primary pathogenic factor is the observation that distal PAVMs are reversible in the Fontan stage, which restores direct hepatopulmonary venous blood flow but not normal pulsatility. However, the proximal collateral vessels do not resolve in the Fontan, pointing to a critical role for pulsatile flow in vascular architecture. In our study, we observe thinning of vSMCs both proximally and distally in the animal model and therefore suggest that pulsatility may be a primary factor in the generation of proximal collateral vessels and an exacerbating factor in the development of distal PAVMs.

The relative contribution of pulsatility and hepatic factor to the pathogenesis of distal AVMs can be assessed by comparing the time of onset in the Glenn and other clinical scenarios. In Glenn patients (no hepatic factor, no pulsatility), AVMs can develop in months (7, 70–72). However, in a rare subset of heterotaxy patients (no hepatic factor, normal pulsatile flow) (73–76), AVMs form more slowly. In these latter patients, who have interrupted inferior vena cava (strongly suggesting left isomerism heterotaxy syndrome (77)), the hepatic veins can (rarely) drain directly to the left atrium in an otherwise normal biventricular heart with intact atrial and ventricular septa. In this scenario (pulsatile pulmonary blood flow, diversion of hepatic venous blood from the lungs), distal PAVMs can take years or even decades to appear (73–76). This suggests that loss of pulsatility critically accelerates the pathogenesis of distal pulmonary AVMs but is not sufficient to induce them.

Thus, while loss of pulsatility may not be the defining factor that leads to distal pulmonary AVMs in the Glenn stage, they are correlated in our rat model. Our immunohistochemical and immunofluorescent evaluation in the Glenn rat model shows dramatic changes of both proximal and distal pulmonary arterial mural cell recruitment Currently, descriptions of dilated distal vessels in the Glenn population exist (78–80); however, there has been no histological analysis of proximal pulmonary arteries. Work in Fontan patients, however, has shown thinning of the proximal pulmonary arterial media (81, 82). The persistence of thin media (proximally, where collaterals form) even as distal PAVMs resolve in the Fontan stage suggests a critical role for pulsatility in maintenance of proximal vascular architecture. We suggest that pulsatility loss is therefore likely a common pathogenic factor in the development of both proximal collateral vessels and distal AVMs.

In summary, we show that the pulsatility of hemodynamic forces in blood vessels is sensed by the endothelium uniquely in each dimension of force. Our data support a model whereby pulsatile stretch induces endothelium to secrete PDGFB, signaling for support from vascular smooth muscle. Indeed, the architecture and cellular composition of the vessel wall change significantly upon loss of pulsatility in the Glenn, with thinning of the vascular wall, thereby shifting the arterial vascular wall structure toward that of a vein, likely impacting downstream endothelium. Together, this work underscores the critical importance of proper blood flow dynamics for the stability of vascular structure and function. In addition, it points to biomechanical signaling pathways as potential novel targets for therapeutic intervention in patients with pulsatility loss, such as in the Glenn circulation. We seek to better understand and address the mechanistic underpinnings of vascular defects that occur during single ventricle palliation and anticipate that our data on pulsatility will provide the foundation for improving clinical outcomes for children that undergo these necessary surgeries.

## Methods

### Data Availability

All sequencing data that support the findings of this study have been deposited in the National Center for Biotechnology Information Gene Expression Omnibus (GEO) and are accessible through the GEO Series accession number GSE298790. All other relevant data are available from the corresponding author on request.

### Combined cardiac catheterization and cardiac MRI

Combined cardiac catheterization and MRI was performed on a Philips XMR setup (Phillips Ingenia Evolution, Best, Netherlands).(83) This consists of a 1.5T MRI-scanner and a BV Pulsera or Allura Clarity or Ingenia (Philips) cardiac X-Ray unit. An appropriate size balloon wedge catheter (Arrow Intl., Reading, PA) and receiver coil were used depending on the weight of the patient.

All studies were conducted under general anesthesia based on clinical need. Patients were recruited and data was recorded in sequence of the patient’s referral for the procedure under University of Texas Southwestern IRB-approved study (STU 032016-009) in 2023 - 2024. All Glenn patients with a single functional ventricle were considered for inclusion in the study. Patients were excluded if they had pulmonary blood flow in addition to the flow provided by the Glenn anastomosis. Both male and female patients were included in the study (see Supplemental Table 1).

Pulsatility was assessed by pulse difference, which was defined as the difference between maximum and minimum value for each parameter of force. Pulsatility index was not selected due to the potential for obscuring differences between patients (see Supplemental Table 2). For each 2D cross-section of the right pulmonary artery obtained, we obtained a mean velocity (averaged over the cross section of the vessel). This data is obtained over 1 to 2 minutes ‘free-breathing’ using phase contrast velocity-encoded cine MRI. Sequence parameters included 40 cardiac phases, TE/TR = 2.7/4.4 ms (TE = echo time, TR = repetition time), with two signal averages, 1.5-2mm × 1.5-2mm × 6-8 mm resolution, SENSE acceleration factor = 2, with the velocity encoding gradient set to 25% above the expected maximum velocity in each vessel. Vendor-provided background phase correction was used. Artificial intelligence-assisted segmentation using Circle (Circle Cardiovascular Imaging, CVI42, v6.1.2) was performed to obtain the maximum and minimum area, from which an idealized circumference was calculated. Pressure tracings were obtained on a standard Siemens Sensis Hemodynamic recording system (Siemens Healthineers, Munich, Germany), using a 5- or 6-French Arrow, Balloon Wedge-Pressure Catheter (Teleflex Medical Headquarter International, Ireland) located in the proximal right branch pulmonary artery.

### In vitro cell-based assays of pulsatile force

Pulsatile force was applied to confluent monolayers of human pulmonary artery endothelial cells (HPAECs, Lonza) for 48 hours under each condition. Pulsatile (1 Hz, 15 dyn/cm^2^) and continuous (15 dyn/cm2) shear was applied using the Ibidi system (Ibidi USA, Wisconsin). Pulsatile (25-5 mm Hg) and continuous (25 mm Hg) pressure was applied using a custom modification to the Ibidi system whereby cells were alternately (1 Hz) exposed to columns of culture media to deliver the desired hydrostatic pressure. Pressure readings were tested and confirmed the using the Siemens Sensis Hemodynamic recording system listed above. Pulsatile stretch was applied using Uniflex plates (1Hz, 1 hr of 3% stretch for acclimation followed by 47 hrs of 10% stretch) as a part of the Flexcell FX-6000 system (Flexcell International, Burlington, NC). Ibidi channels were pre-coated with the proprietary Ibi-treat to promote cell adhesion, and Flexcell uniflex plates were coated with 0.1% gelatin to promote cell adhesion.

All experiments utilized at least three biological replicates, using separate aliquots of HPAECs from two different donors. RNA was extracted using RNeasy Plus kit (Qiagen) prior to bulk RNA sequencing. Library generation and bulk RNA sequencing was carried out by the McDermott Next Generation Sequencing core facility at UT Southwestern Medical Center, using an Illumina NextSeq 2000. Processed sequencing data (using iGenomes annotations) was provided by the core, and a TPM cutoff of 10 was applied (if any test condition resulted in a gene’s TPM >10, the gene was included in our analysis). PCA plots of each condition are shown in **Supplementary Figure 4**.

For pathway analysis, we performed gene set enrichment analysis was performed for sets curated in MsigDb (PMID 26771021) using with fgsea (BioRxiv https://doi.org/10.1101/060012) Bioconductor package. To simultaneously account for both the magnitude and significance of the measured effects, genes were ranked by the quotient of log fold-change and adjusted P-value.

### Human Lung Single Cell RNA sequencing

The LungMAP Human Lung CellRef atlas was utilized via the online portal at www.lungmap.net. (38, 39) This dataset includes 148 normal human lung samples from 104 donors, including child, adolescent, and adult tissue. Tissue types include parenchyma, trachea, bronchi, bronchus, and small airways. 347,970 cells from the dataset were included in our analysis. Both ShinyCell and cell x gene were utilized for visualization of cell-specific gene expression. Dataset ID: LMEX0000004396.

### Measurement of PDGFB secretion in vitro

Cultured HPAECs, human arterial endothelial cells (HAECs, Lonza), human pulmonary artery smooth muscle cells (PASMCs, Lonza), and normal human lung fibroblasts (NHLFs, Lonza were plated onto Flexcell uniflex plates (pre-coated with collagen I). Each cell line was grown in the recommended cell culture media, obtained from Lonza: HPAECs and HAECs were grown in EGM-2, PASMCs in SmGM-2, and NHLFs in FGM-2. Cells were grown to confluency then subjected to pulsatile stretch for 48 hours as above. Media was collected and PDGF-BB was measured by ELISA (R+D systems DY220). For comparison to endothelial cells under flow, media was obtained from confluent HPAECs in 6 well plates under continuous flow at 15 dyn/cm^2^ shear stress in a rotational flow apparatus (Thermo Scientific Model No: 88881101).

For measurement of transpulmonary gradient of PDGF-BB (here termed PDGFB) in human plasma, samples were obtained directly from Glenn and heart transplant patients in the course of regularly indicated cardiac catheterization. Patients were consented for the IRB approved study (University of Texas Southwestern IRB: UTSW-2020-0047). Samples were obtained from the pre-capillary blood (right pulmonary artery) and the post-capillary blood (right pulmonary wedge position, with the catheter in the distal pulmonary artery with balloon inflated and the sample drawn through the end hole) in each patient. Nature of each sample as pre- or post-capillary was then confirmed by prompt measurement of oxygen saturation, and only fully oxygenated samples were used for the analysis. All samples were obtained in K2-EDTA purple-top tubes, placed on ice, and promptly spun down for plasma isolation. Samples were stored at −80°C up to 2 years maximum prior to measurement by ELISA as above.

### Surgical Rat Model of Glenn Circulation

The rat surgical model of Glenn circulation is described in detail previously.(36) Briefly, we performed an end-to-end cavopulmonary anastomosis between the left superior vena cava (L-SVC) and the left pulmonary artery (LPA). This was previously shown to result in progressive impairment (beginning by two weeks) in oxygenation in tandem with the development of intrapulmonary arteriovenous shunts as detected by both bubble echocardiography and microsphere injection.

Adult Sprague–Dawley rats (male and female, 6–12 weeks of age) were used for all experiments (Taconic). All rats were housed in the Biomedical Research Center at the Medical College of Wisconsin with access to standard chow diet (PicoLab Lab Diet, 5L0D), water ad libitum, and maintained in a 12-hour light/dark cycle. All surgical procedures were performed under isoflurane anesthesia (1%–3%). All experimental protocols were approved by the Medical College of Wisconsin Institutional Animal Care and Use Committee prior to initiation of experimental protocols (Animal Use Agreement #7731).

### Tissue Preparation and Immunohistochemistry (IHC) /Immunofluorescence (IF)

Lungs were harvested at 2 and 6 months after surgery for these experiments. Lungs were inflated to 10 mmHg for 10 minutes with 10% neutral buffered formalin to promote normal alveolar architecture in sections, followed by overnight fixation in 10% neutral buffered formalin. Formalin-fixed, paraffin-embedded sections were prepared using the left lung of the Glenn and sham operated animals. IHC staining was done for SMA (Abcam, ab7817) using the Leica Bond Rx automated staining platform with Leica BOND Polymer Refine Detection kit (Leica, DS9800) per manufacturer instructions, using standard labeled-streptavidin-biotin detection with secondary antibody (Jackson Immuno Research Labs, Biotinylated donkey anti-mouse 1:500, #715-066-151), streptavidin HRP (Vector Laboratories, #SA-5004), and DAB chromogen (BioCare, BDB2004).

Slides were scanned using Hamamatsu HT whole slide scanner (Hamamatsu, USA) and analyzed with QuPath (Edinburgh, V0.5.1). The file names were randomized and blinded prior to analysis. Due to the possibility of changes in vessel size between sham and Glenn-operated rates, measurements were taken from the most proximal pulmonary arteries and veins to ensure a valid comparison between samples. Arteries were differentiated from veins in consultation with a trained pathologist, by their classic histological characteristics and anatomical proximity to the mainstem bronchi. Slides were not utilized for measurement if no major bronchi or large vessels could be identified. Measurements were taken only in areas where the thickness was uniform and the endothelial cells formed a clear monolayer, to avoid artifactual error from the angle of the slice.10 measurements of each large vessel (artery or vein, Glenn or sham-operated) were obtained, and the mean measurement value was used for analysis.

We used IF staining and confocal microscopy to identify distal vessels for quantification of endothelial PDGFB protein and perivascular smooth muscle. Lung sections were prepared as above. Sections were permeabilized for 10 min in 0.2% Triton X-100 in PBS, then subjected to antigen retrieval with R-buffer B in a 2100 Retriever (Electron Microscopy Sciences). Slides were blocked with CAS-block (Invitrogen), and primary antibody incubations were carried out at 4°C overnight. Primary antibodies were as follows: VE-cadherin/CDH5 (R+D systems, AF1002, 1:100), PDGFB (Santa Cruz, sc-365805, 1:100), and SMA-APC (Abcam, ab225143, 1:400). Imaging was performed on a Nikon CSU-W1 dual-camera inverted spinning-disc confocal microscope. Vessels were chosen as <50 µm and >20 µm in diameter, with a clear lumen and apical VE-cadherin staining visible. Images were analyzed in Fiji as follows: the percentage of the circumference of each vessel covered by SMA staining was calculated, and a mask was made of the pixels positive for SMA staining in the sub-endothelial region and the mean pixel intensity of those pixels was calculated.

### Statistical Analysis

Sample sizes were based on patient availability and standards in the field. Data were both analyzed and plotted in Graphpad PRISM 10.2.3. Descriptive statistics (mean, standard deviation, median, min, and max) were employed for summarizing demographic, hemodynamic, and clinical variables. Statistical tests (indicated in the relevant methods sections and figure legends) utilized were the parametric unpaired Student’s *t*-test and the one-way ANOVA with Tukey’s multiple comparisons test. Statistical significance is noted as follows: ns = *p* > 0.05; * = 0.01 < *p* < 0.05; ** = 0.001 < *p* < 0.01; *** = 0.0001 < *p* < 0.001; **** = *p* < 0.0001. Superplots were made with individual colors representing biological replicates (individual rats), and the mean of the averages of each biological replicate is shown (84). Cell images were analyzed in FIJI. Figures and models were made using Microsoft PowerPoint, data were partially analyzed in Microsoft Excel, and text was written in Microsoft Word.

## ACKNOWLEDGEMENTS

We are grateful to the entire Cleaver Lab for useful discussions and reading of the manuscript. We thank Denise Marciano for the use of her lab’s Flexcell apparatus. We thank the UT Southwestern Medical Center (UTSW) Quantitative Light Microscopy Core Facility (Marcel Mettlen, Ph.D.) for training/assistance. The results here are in part based upon data generated by the LungMAP Consortium and downloaded from (www.lungmap.net), on March 15, 2025. The LungMAP consortium, the Human Tissue Core (U01-HL144861), and the LungMAP Data Coordinating Center (U24-HL148865) are funded by the National Heart, Lung, and Blood Institute (NHLBI).

## Author contributions

Conceptualization: S.S., T.H., M.F., M.L.I.A., O.C.; Methodology: S.S., T.H., M.F., S.R., A.D.S., O.C.; Investigation: S.S., L.T., T.W., S.R., O.C.; Resources: S.S., T.W., A.D.S., T.C., O.C.; Data curation: S.S., M.A.C., L.T., T.H.; Writing - original draft: S.S., O.C.; Writing - review & editing: S.S., T.H., M.L.I.A., A.D.S., O.C.; Visualization: M.A.C., S.S.; Supervision: S.S., O.C.; Project administration: S.S., O.C.; Funding acquisition: S.S., T.C., A.D.S., O.C.

## Non-standard Abbreviations and Acronyms

AV: arteriovenous
GSEA: Gene Set Enrichment Analysis
HAECs: human aortic endothelial cells
HPAECs: human pulmonary arterial endothelial cells
IF: immunofluorescence
IHC: immunohistochemistry
LPA: left pulmonary artery
L-SVC: left superior vena cava
NHLFs: normal human lung fibroblasts
PA: pulmonary artery
PASMCs: human pulmonary artery smooth muscle cells
PAVMs: microvascular pulmonary arteriovenous malformations
PDGFB: platelet derived growth factor B
PDGFR: platelet derived growth receptor
RPA: right pulmonary artery
SVC: superior vena cava
SV-CHD: Single ventricle congenital heart disease
vSMC: vascular smooth muscle cell

