## Supplemental Figures for "Pulsatile flow dynamics maintain pulmonary arterial architecture"

**Supplemental Fig 1: Comparison of arterial and venous characteristics highlights the definitional importance of pulsatility**

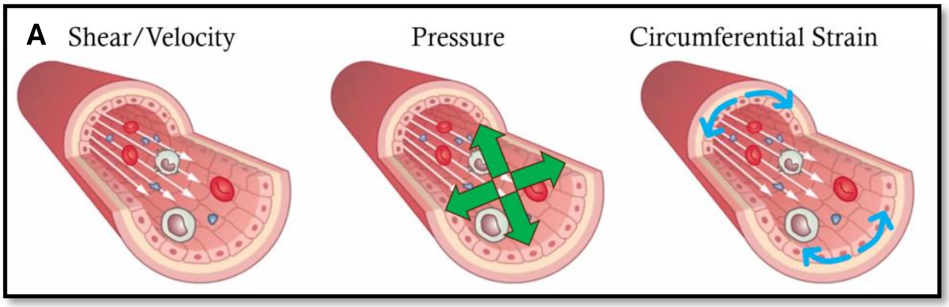


| B | | Systemic | | Pulmonary | | Umbilical | |
|---|---|---|---|---|---|---|---|
| | | Artery | Vein | Artery | Vein | Artery | Vein |
| Hemodynamic Forces | Pulsatility | YES | NO | YES | NO | YES | NO |
| | Mean Pressure (mm Hg) | 70 | 4 | 12 | 6 | 50 | 20 |
| | Peak Velocity (cm/s) | 100 | 15 | 75 | 40 | 45 | 15 |
| Gas Transport | PO2 | 100 | 40 | 40 | 100 | 20 | 30 |
| | PCO2 | 40 | 45 | 45 | 40 | 50 | 40 |
| | pH | 7.4 | 7.36 | 7.36 | 7.4 | 7.27 | 7.35 |
| | Hgb Saturation | 100% | 70% | 70% | 100% | 50% | 70% |
| | Hgb Isoforms | HgbA/B | HgbA/B | HgbA/B | HgbA/B | HgbF | HgbF |
| Histological Features | Intima | + | + | + | + | + | + |
| | Internal Elastic Lamina | ++ | + | ++ | + | - | - |
| | Media (VSMCs) | +++ | + | +++ | + | +++ | + |
| | Adventitia | ++ | + | ++ | + | ++ | + |
| | Forms Valves | NO | YES | NO | NO | NO | NO |

**Supplemental Fig 1: Comparison of arterial and venous characteristics highlights the definitional importance of pulsatility. a** Three dimensions of hemodynamic force on endothelial cells. b Comparison of key features of the three major anatomical sets of arteries and veins. Pulsatility is common to all arteries despite their pressure, oxygenation, or peak shear stress/velocity.

### Supplemental Figure 2: Comparison of baseline transcriptome of key endothelial genes under static conditions

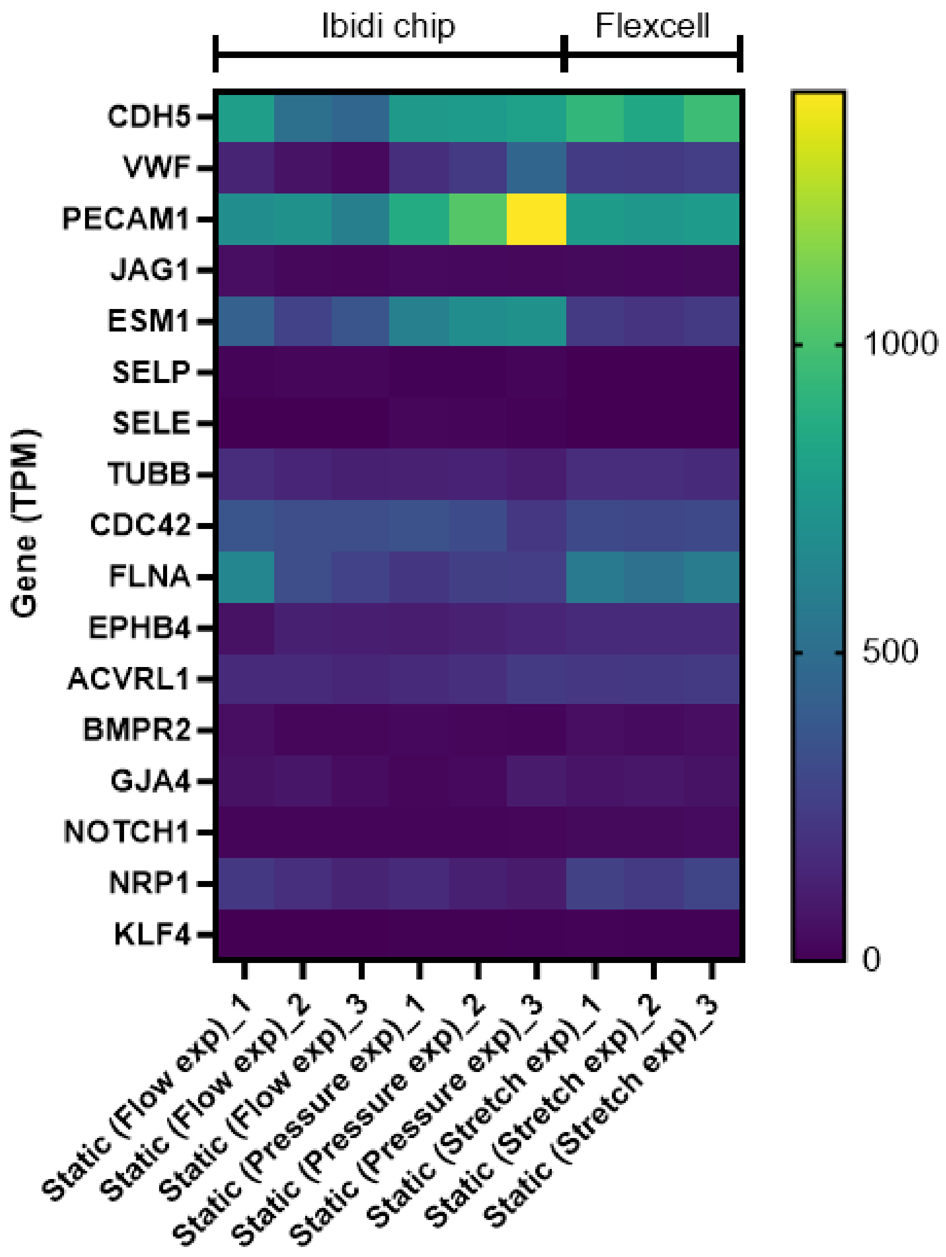


**Supplemental Figure 2: Comparison of baseline transcriptome of key endothelial genes under static conditions.** Heatmap comparing baseline expression of key endothelial cell genes shows that despite being seeded on different substrates, endothelial cells used in our experiments did not vary in their baseline conditions prior to the application of pulsatile or non-pulsatile force.

### Supplemental Figure 3: Impact of pulsatility on significantly regulated genes

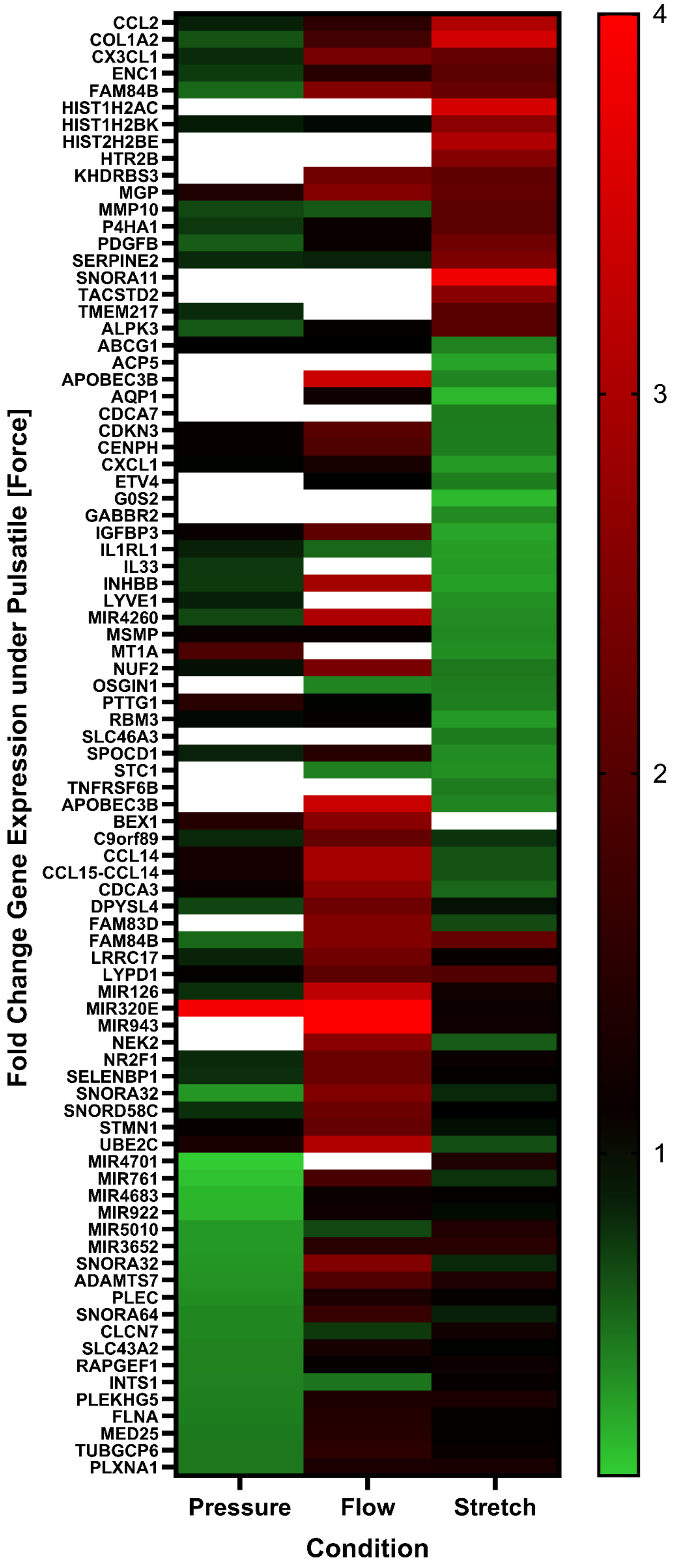


**Supplemental Figure 3: Impact of pulsatility on significantly regulated genes.** (Alternate visualization of data in figure 3) Heatmap showing fold change in gene expression under different dimensions of force, comparing pulsatile to non-pulsatile conditions. White boxes indicate gene was excluded from analysis for that condition due to low expression (TPM < 10 for that gene under both pulsatile and non-pulsatile conditions).

**Supplemental Fig 4: PCA plots of bulk RNAseq data**

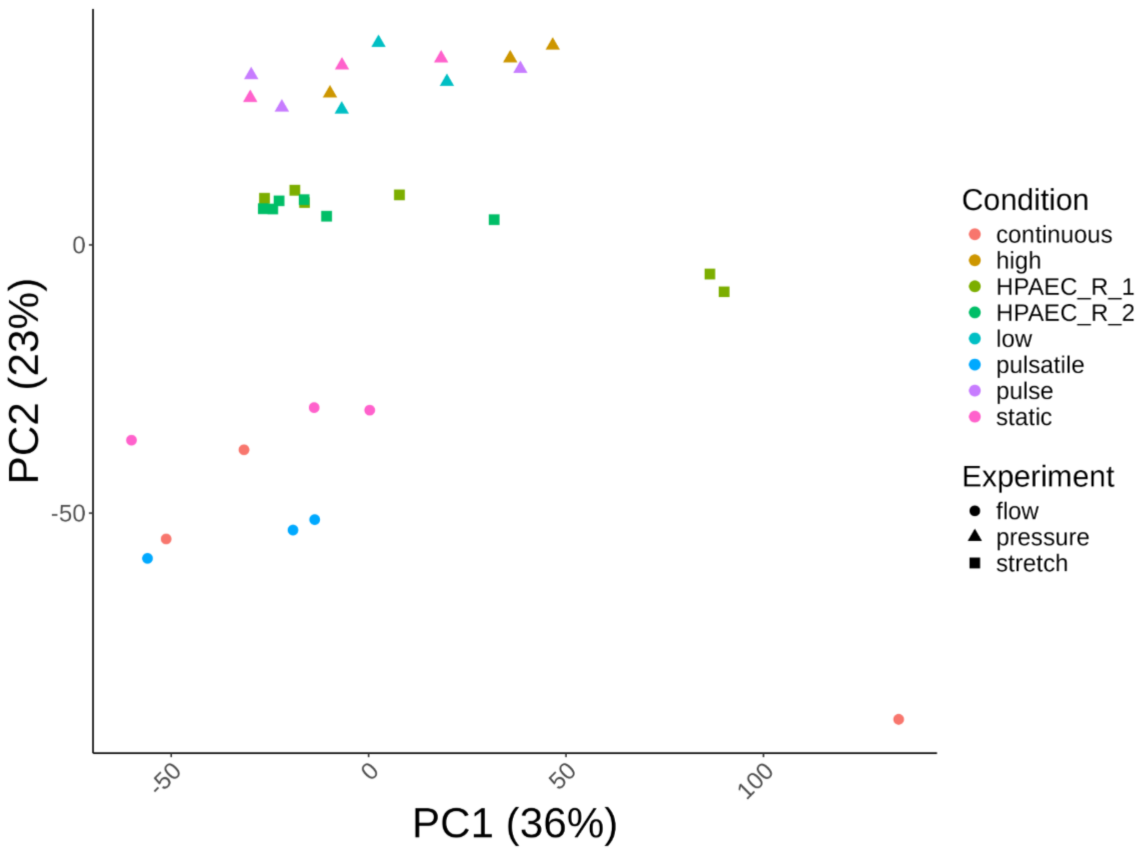


**Supplemental Figure 4: PCA plots of bulk RNAseq data**
Principal component analysis of the bulk RNA sequencing data used in the analysis. Percentages on axes denote percent of variation accounted for by each principal component. HPAEC_R_1 and _2 denotes no stretch and pulsatile stretch conditions, respectively.

#### Supplemental Table 1: Patient Demographics

| | Glenn | Controls |
|---|---|---|
| *Demographics* | n = 20 | n = 17 |
| Age in years | 4.4 (1.5) | 4.5 (0.9) |
| Male | 12 (60%) | 8 (90%) |
| *Cardiac Diagnosis* | | |
| Single ventricle - Glenn stage | 20 (100%) | |
| Systemic left ventricle | 9 (45%) | 17 (100%) |
| Heart Transplantation | | 9 |
| Atrial septal defect | | 4 |
| Coarctation of the aorta | | 2 |
| Bicuspid aortic valve | | 1 |
| Ebsteins Anomaly | | 1 |
| *Hemodynamic State* | | |
| Cardiac Index | 4.1 (0.5) | 2.8 (0.5) |

**Supplemental Table 1: Patient Demographics.** Descriptive statistics of patient data included in the study. Data is presented as mean (standard deviation) unless otherwise noted.

### Supplemental Table 2: Comparison of Pulsatility Index and Pulse Difference

| max | min | mean | PD | PI |
|---|---|---|---|---|
| 20 | 5 | 10 | **15** | **1.5** |
| 30 | 5 | 16.2 | **25** | **1.5** |
| 40 | 10 | 20 | **30** | **1.5** |
| 50 | 15 | 23 | **35** | **1.5** |

**Supplemental Table 2: Comparison of Pulsatility Index and Pulse Difference.** Comparison of pulsatility index (PI) and pulse difference (PD) shows that PI can be unchanged for a variety of different PD.
